## Supplementary figures and images for "Deriving time-concordant event cascades from gene expression data: A case study for Drug-Induced Liver Injury (DILI)"

### S1_Figure

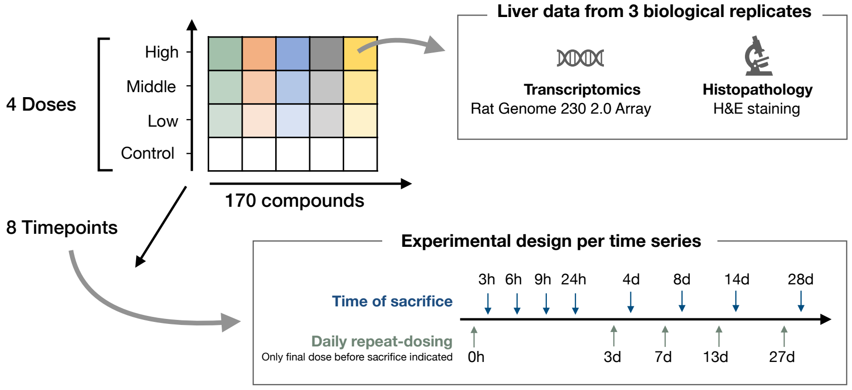

### S2_Figure

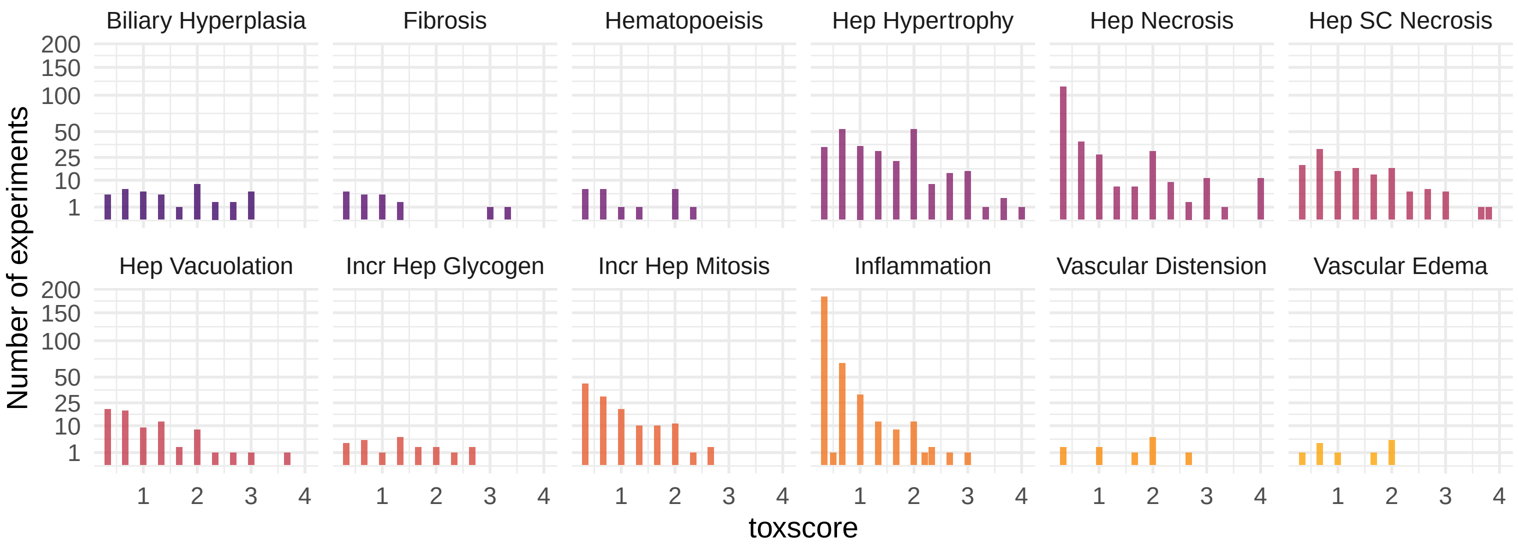

### S3_Figure

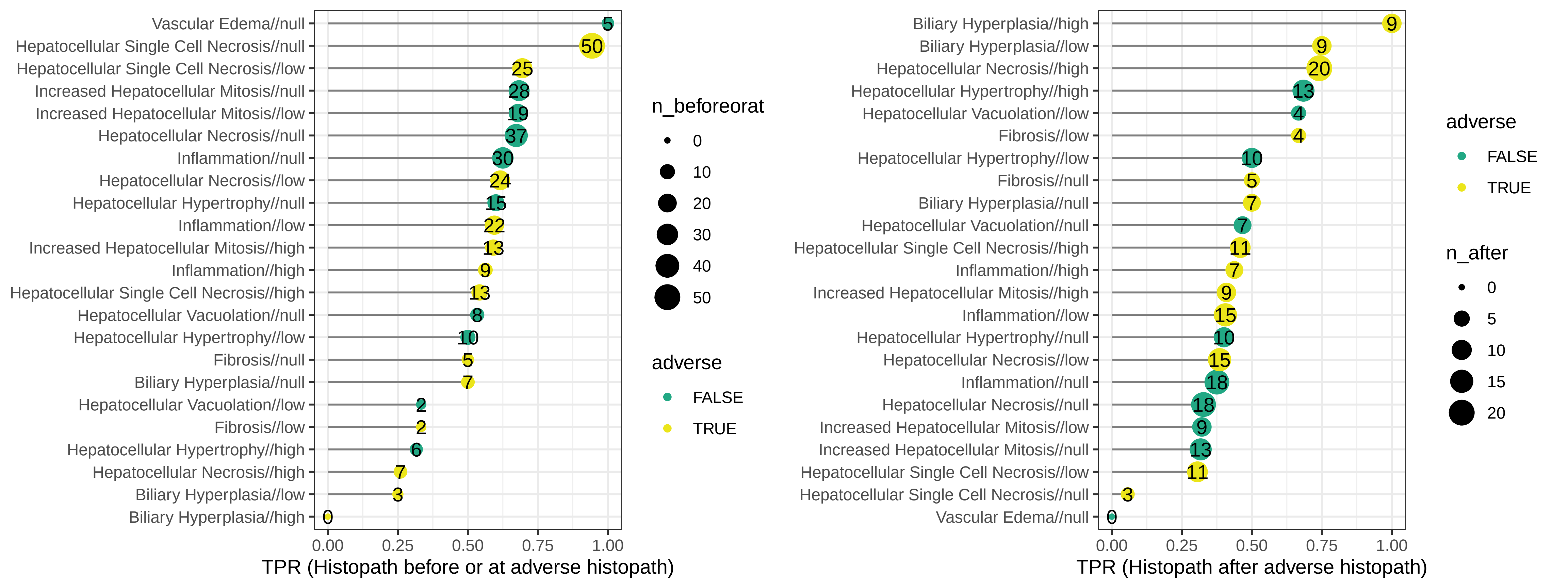

### S4_Figure

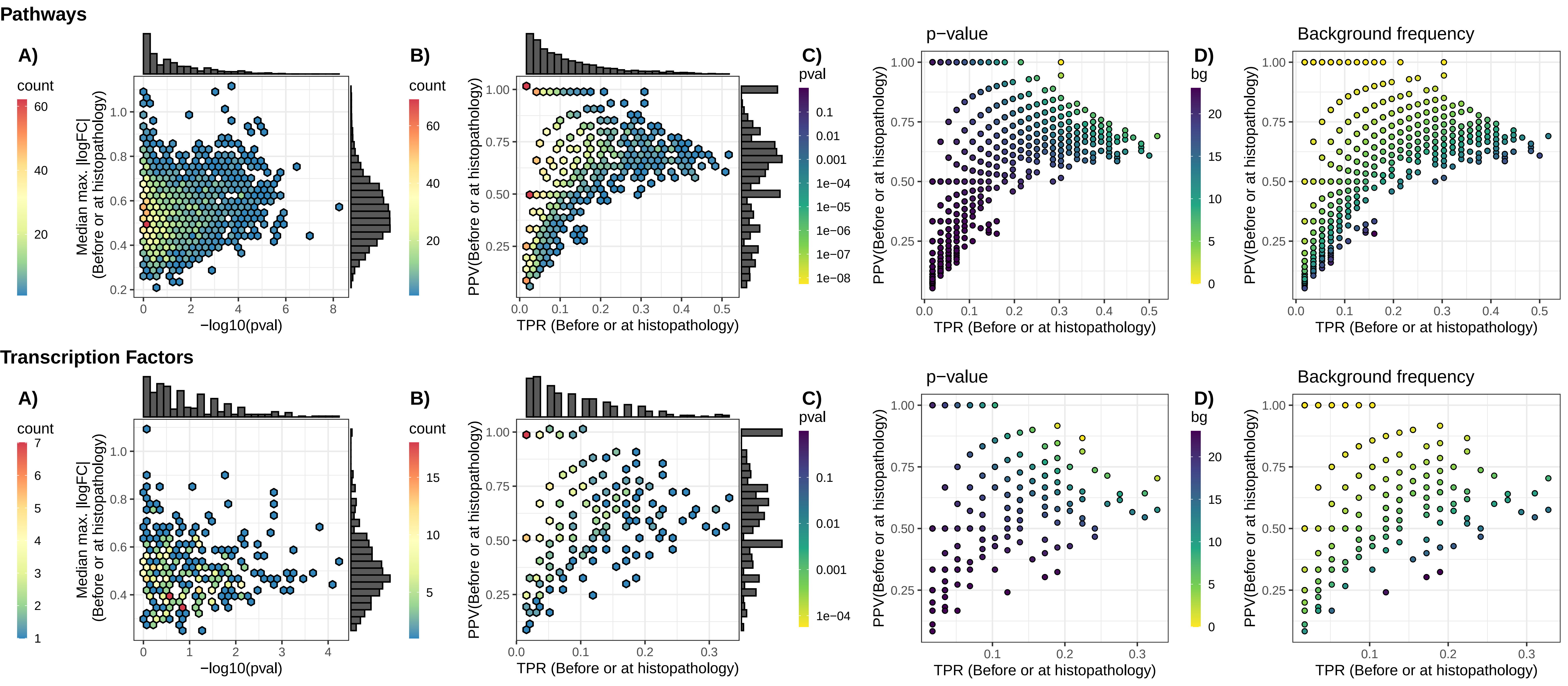
