## Supplementary material for "Deriving time-concordant event cascades from gene expression data: A case study for Drug-Induced Liver Injury (DILI)": S1_Table

|  |  | cpAOP | Probablistic qAOP | Mechanistic qAOP | This study |
| --- | --- | --- | --- | --- | --- |
| Output KERs | Coverage of events | All events with known associations  (++) | Limited to AOP scaffold  (-) | Limited to AOP scaffold  (-) | All events in data  (+) |
|  | Evidence for causality | No | No | Yes (Experimental) | Yes (Time concordance) |
|  | Evidence for predictivity | No | Yes | No | Yes (PPV) |
| Purpose | Support AOP development | Yes (Potentially new KER) | No (Uses existing AOP) | No (Uses existing AOP) | Yes (Potentially new KER) |
|  | Make AOP useful for risk assessment? | No (Does not add additional info) | Yes (Enables prediction) | Yes (Better understanding) | Yes (Better understanding) |
| Approach | Automatable? | Yes | Yes | No (New experiments needed) | Yes |
|  | Large number of chemicals | Yes (All compounds with known associations) | Yes (All compounds with relevant assay data) | No | Yes (All compounds with time-series data) |
|  | Based on in vivo data | No (potentially indirectly) | No | Yes | Yes |
|  | Transferrable to other AE? | Yes | Yes (if AOP available) | Partially (New experiments needed) | Partially (if suitable time series data is available) |
