## Supplementary material for "Deriving time-concordant event cascades from gene expression data: A case study for Drug-Induced Liver Injury (DILI)": S2_Table

|  | **COMPOUND_NAME** | **DILIst Classification** | **vDILIConcern** | **Severity.Class** | **29 day** | **4 day** |
| --- | --- | --- | --- | --- | --- | --- |
| **1** | acetamidofluorene |  |  |  | Middle, High |  |
| **2** | acetaminophen | 1 | vMost-DILI-Concern | 5 |  |  |
| **3** | aspirin | 1 | vLess-DILI-Concern | 0 |  |  |
| **4** | bendazac | 1 | vMost-DILI-Concern | 8 | High | High |
| **5** | bromobenzene | 1 |  |  |  |  |
| **6** | captopril | 1 | vLess-DILI-Concern | 7 |  |  |
| **7** | clofibrate | 1 | vLess-DILI-Concern | 3 | Middle, High |  |
| **8** | clomipramine | 1 | vMost-DILI-Concern | 8 | High |  |
| **9** | colchicine | 0 | Ambiguous DILI-concern | 6 |  |  |
| **10** | coumarin |  |  |  | High |  |
| **11** | danazol | 1 | vMost-DILI-Concern | 8 |  |  |
| **12** | dantrolene | 1 | vMost-DILI-Concern | 8 |  |  |
| **13** | diclofenac | 1 | vMost-DILI-Concern | 8 |  |  |
| **14** | ethambutol | 1 | vMost-DILI-Concern | 8 | Middle, High |  |
| **15** | ethinylestradiol | 1 |  |  |  |  |
| **16** | ethionamide | 1 | vLess-DILI-Concern | 3 | High | High |
| **17** | ethionine |  |  |  |  |  |
| **18** | fenofibrate | 1 | vLess-DILI-Concern | 3 | Low, Middle, High | Middle, High |
| **19** | gemfibrozil | 1 | vMost-DILI-Concern | 4 | Low, Middle, High |  |
| **20** | ibuprofen | 1 | vLess-DILI-Concern | 3 |  |  |
| **21** | ketoconazole | 1 | vMost-DILI-Concern | 8 |  |  |
| **22** | lomustine | 1 | vLess-DILI-Concern | 3 | High |  |
| **23** | methapyrilene | 1 |  |  | Middle, High | High |
| **24** | methyltestosterone | 1 | vLess-DILI-Concern | 2 |  |  |
| **25** | monocrotaline |  |  |  | Middle, High |  |
| **26** | nitrosodiethylamine |  |  |  | Middle, High | High |
| **27** | phalloidin |  |  |  | High | High |
| **28** | phenacetin | 1 |  |  |  |  |
| **29** | simvastatin | 1 | vLess-DILI-Concern | 3 |  |  |
| **30** | theophylline | 0 | vNo-DILI-Concern | 0 | High |  |
| **31** | thioacetamide |  |  |  | Low, Middle, High | High |
| **32** | WY-14643 |  |  |  | Low, Middle, High | Low, Middle, High |
