## Supplementary material for "Deriving time-concordant event cascades from gene expression data: A case study for Drug-Induced Liver Injury (DILI)": S4_Table

| Event | Direction | pval | TPR | PPV | logFC |
| --- | --- | --- | --- | --- | --- |
| **Reactome pathways** | | | | | |
| Dectin-1 mediated noncanonical NF-kB signaling | Up | 8.61E-05 | 0.446 (25/56) | 0.625 (25/40) | 0.79 |
| Recycling of eIF2:GDP | Up | 9.53E-06 | 0.446 (25/56) | 0.676 (25/37) | 0.94 |
| RNA Polymerase I Promoter Escape | Up | 1.67E-06 | 0.446 (25/56) | 0.714 (25/35) | 0.61 |
| RNA Polymerase I Transcription Termination | Up | 4.12E-06 | 0.446 (25/56) | 0.694 (25/36) | 0.67 |
| rRNA modification in the nucleus and cytosol | Up | 8.61E-05 | 0.446 (25/56) | 0.625 (25/40) | 0.86 |
| SCF(Skp2)-mediated degradation of p27/p21 | Up | 8.61E-05 | 0.446 (25/56) | 0.625 (25/40) | 0.78 |
| tRNA modification in the nucleus and cytosol | Up | 3.65E-06 | 0.464 (26/56) | 0.684 (26/38) | 0.6 |
| Folding of actin by CCT/TriC | Up | 1.48E-05 | 0.482 (27/56) | 0.643 (27/42) | 0.96 |
| Nonsense-Mediated Decay (NMD) | Up | 6.98E-06 | 0.482 (27/56) | 0.659 (27/41) | 0.73 |
| Response of EIF2AK4 (GCN2) to amino acid deficiency | Up | 6.98E-06 | 0.482 (27/56) | 0.659 (27/41) | 0.74 |
| rRNA processing | Up | 2.99E-05 | 0.482 (27/56) | 0.628 (27/43) | 0.75 |
| Activation of the mRNA upon binding of the cap-binding complex and eIFs, and subsequent binding to 43S | Up | 4.69E-05 | 0.5 (28/56) | 0.609 (28/46) | 0.8 |
| Cytosolic tRNA aminoacylation | Up | 4.11E-07 | 0.518 (29/56) | 0.69 (29/42) | 0.74 |
| Mitotic G1 phase and G1/S transition | Up | 1.79E-06 | 0.393 (22/56) | 0.759 (22/29) | 0.5 |
| Defects in biotin (Btn) metabolism | Down | 1.67E-06 | 0.375 (21/56) | 0.778 (21/27) | -0.73 |
| HS-GAG biosynthesis | Down | 1.58E-06 | 0.286 (16/56) | 0.889 (16/18) | -0.56 |
| DNA Replication | Up | 1.46E-06 | 0.357 (20/56) | 0.8 (20/25) | 0.55 |
| S Phase | Up | 1.46E-06 | 0.357 (20/56) | 0.8 (20/25) | 0.52 |
| Cell Cycle Checkpoints | Up | 1.18E-06 | 0.339 (19/56) | 0.826 (19/23) | 0.5 |
| Diseases associated with glycosaminoglycan metabolism | Down | 7.44E-08 | 0.304 (17/56) | 0.944 (17/18) | -0.45 |
| Recycling of bile acids and salts | Down | 5.66E-09 | 0.304 (17/56) | 1 (17/17) | -0.57 |
| Tyrosine catabolism | Down | 7.23E-06 | 0.286 (16/56) | 0.842 (16/19) | -0.91 |
| Beta-oxidation of very long chain fatty acids | Up | 3.42E-05 | 0.339 (19/56) | 0.731 (19/26) | 0.92 |
| Removal of aminoterminal propeptides from gamma-carboxylated proteins | Down | 3.42E-05 | 0.339 (19/56) | 0.731 (19/26) | -0.92 |
| Alpha-oxidation of phytanate | Up | 1.99E-04 | 0.268 (15/56) | 0.75 (15/20) | 0.96 |
| Cholesterol biosynthesis | Down | 9.83E-03 | 0.304 (17/56) | 0.567 (17/30) | -0.99 |
| mitochondrial fatty acid beta-oxidation of saturated fatty acids | Up | 4.50E-04 | 0.232 (13/56) | 0.765 (13/17) | 1.02 |
| Beta oxidation of decanoyl-CoA to octanoyl-CoA-CoA | Up | 1.09E-03 | 0.214 (12/56) | 0.75 (12/16) | 1.08 |
| mitochondrial fatty acid beta-oxidation of unsaturated fatty acids | Up | 1.75E-04 | 0.357 (20/56) | 0.667 (20/30) | 1.11 |
| **Transcription Factors** | | | | | |
| Etv4 | Down | 7.51E-04 | 0.259 (15/58) | 0.714 (15/21) | -0.5 |
| Zfp217 | Down | 1.10E-02 | 0.259 (15/58) | 0.6 (15/25) | -0.58 |
| Kdm5b | Down | 5.73E-03 | 0.276 (16/58) | 0.615 (16/26) | -0.62 |
| Zbtb11 | Up | 3.13E-03 | 0.276 (16/58) | 0.64 (16/25) | 0.57 |
| Nfe2l2 | Up | 1.42E-02 | 0.293 (17/58) | 0.567 (17/30) | 0.79 |
| Atf4 | Up | 1.41E-03 | 0.31 (18/58) | 0.643 (18/28) | 0.84 |
| Srebf1 | Down | 1.41E-03 | 0.31 (18/58) | 0.643 (18/28) | -0.74 |
| Srebf2 | Down | 1.93E-02 | 0.31 (18/58) | 0.545 (18/33) | -0.9 |
| E2f2 | Up | 1.44E-04 | 0.328 (19/58) | 0.704 (19/27) | 0.69 |
| Nr1h3 | Down | 6.84E-03 | 0.328 (19/58) | 0.576 (19/33) | -0.66 |
| Ebf1 | Down | 7.15E-04 | 0.241 (14/58) | 0.737 (14/19) | -0.43 |
| Foxl2 | Up | 6.62E-04 | 0.155 (9/58) | 0.9 (9/10) | 0.59 |
| Hnf4a | Down | 6.62E-04 | 0.155 (9/58) | 0.9 (9/10) | -0.49 |
| Mafb | Down | 6.62E-04 | 0.155 (9/58) | 0.9 (9/10) | -0.48 |
| Tead1 | Down | 4.03E-04 | 0.19 (11/58) | 0.846 (11/13) | -0.47 |
| Sox13 | Down | 2.17E-04 | 0.224 (13/58) | 0.812 (13/16) | -0.5 |
| Hoxb13 | Down | 8.86E-05 | 0.19 (11/58) | 0.917 (11/12) | -0.45 |
| Tfap4 | Down | 5.77E-05 | 0.224 (13/58) | 0.867 (13/15) | -0.53 |
| Pou2f2 | Down | 4.03E-02 | 0.207 (12/58) | 0.571 (12/21) | -0.65 |
| Tfap2c | Down | 3.13E-02 | 0.155 (9/58) | 0.643 (9/14) | -0.7 |
| Irf9 | Up | 4.03E-02 | 0.207 (12/58) | 0.571 (12/21) | 0.74 |
| Klf4 | Up | 1.31E-03 | 0.19 (11/58) | 0.786 (11/14) | 0.79 |
