## Supplementary material for "Deriving time-concordant event cascades from gene expression data: A case study for Drug-Induced Liver Injury (DILI)": S5_Table

| class | Preceding event | Later event | TPR (Before or at) | TPR (Before) | Sources |
| --- | --- | --- | --- | --- | --- |
| PPI | Mef2c(Down) | Myod1(Down) | 0.667 (4/6) | 0.333 (2/6) | BioGRID;Lit-BM-17;SIGNOR;Wang |
|  | Nr1h2(Down) | Ppara(Down) | 0.3 (3/10) | 0.1 (1/10) | SignaLink3 |
|  | Pax6(Down) | Maf(Down) | 0.333 (3/9) | 0.111 (1/9) | SPIKE |
|  | Ppara(Down) | Nr1h2(Down) | 0.444 (4/9) | 0.222 (2/9) | SignaLink3 |
| Regulon | Cebpa(Down) | Hnf4a(Down) | 0.333 (3/9) | 0 (0/9) | A |
|  | Elf3(Down) | Meis1(Down) | 0.364 (4/11) | 0 (0/11) | C |
|  | Hnf1a(Down) | Hnf4a(Down) | 0.667 (6/9) | 0 (0/9) | A |
|  | Hnf4a(Down) | Cebpa(Down) | 0.75 (3/4) | 0 (0/4) | A |
|  | Nfe2l1(Down) | Tead1(Down) | 0.727 (8/11) | 0 (0/11) | C |
|  | Nr1h2(Down) | Srebf1(Down) | 0.222 (4/18) | 0 (0/18) | C |
|  | Nr1h3(Down) | Srebf1(Down) | 0.5 (9/18) | 0.278 (5/18) | C |
|  | Pbx2(Down) | Meis1(Down) | 0.636 (7/11) | 0 (0/11) | C |
|  | Pbx3(Down) | Meis2(Down) | 0.636 (7/11) | 0 (0/11) | C |
|  | Pdx1(Down) | Hnf4a(Down) | 0.667 (6/9) | 0.444 (4/9) | C |
|  | Prdm1(Down) | Tead1(Down) | 0.364 (4/11) | 0 (0/11) | C |
|  | Rara(Down) | Hnf4a(Down) | 0.222 (2/9) | 0 (0/9) | A |
|  | Rela(Up) | Nfkb1(Up) | 1 (4/4) | 0 (0/4) | A |
|  | Sox11(Down) | Tead1(Down) | 0.273 (3/11) | 0 (0/11) | C |
|  | Tal1(Down) | Nfkb1(Up) | 0.25 (1/4) | 0 (0/4) | A |
|  | Tcf12(Down) | Tead1(Down) | 0.818 (9/11) | 0 (0/11) | C |
|  | Tcf4(Down) | Tead1(Down) | 0.545 (6/11) | 0 (0/11) | C |
|  | Zfp384(Down) | Meis2(Down) | 0.636 (7/11) | 0 (0/11) | C |
|  | Zfx(Up) | Zfx(Up) | 1 (10/10) | 0 (0/10) | E |
